# Chronic therapy with α1A-adrenergic agonist reverses RV failure and mitochondrial dysfunction

**DOI:** 10.64898/2026.03.18.712768

**Authors:** OY Li, PM Swigart, N Reddy, BE Myagmar, E Bat-Erdene, PC Simpson, AJ Baker

**Author notes:** Correspondence to Anthony J. Baker, Ph.D., University of California, San Francisco, VA Medical Center, Cardiology Division (111C), 4150 Clement St, San Francisco, CA 94121.

## Abstract

Right ventricular failure (RVF) is a serious disease with a high mortality but no effective pharmacologic treatments. We reported RVF was reversed by chronic treatment with an α1A-adrenergic receptor (α1A-AR) agonist. Recent studies suggest mitochondrial dysfunction contributes to RVF. Therefore, we investigated if reversal of RVF by chronic α1A-AR agonist treatment involved improved mitochondrial function. A mouse model of RVF caused by pulmonary artery constriction (PAC) for 2 wk was chronically treated for a further 2 wk. with a low dose of the α1A-AR agonist A61603 (10 ng/kg/day) or vehicle (no drug control). RV dysfunction was assessed from the fractional shortening of the RV outflow tract (RVOT FS). RVOT FS for sham controls (46.5 ± 1.3 %, n = 9) was reduced 4 wk after PAC (27.6 ± 1.5 %, n = 13, *P* < 0.0001), but was higher after PAC plus 2 wk A61603 treatment (34.5 ± 0.6 %, n = 14, *P* < 0.001). RV myocardial respiration rate (O_2_ consumption) for sham controls (776 ± 51 pM/s/mg, n = 9) was reduced 4 wk after PAC (493 ± 28 pM/s/mg, n = 15, *P* <0.0001), but was higher after PAC plus 2 wk A61603 treatment (634 ± 30 pM/s/mg, n = 11, *P* <0.05). RV myocardial ATP level for sham controls (3.3 ± 0.1 mM, n = 10) was reduced 4 wk after PAC (1.9 ± 0.1 mM, n = 6, *P* < 0.0001), but was higher after PAC plus 2 wk A61603 treatment (2.6 ± 0.13 mM, n = 7, *P* < 0.01). In conclusion, reversal of RVF after chronic A61603 treatment involved reversal of mitochondrial dysfunction. Consistent with our previous studies, this study suggests that the α1A-AR is a therapeutic target to treat RVF.

**Highlights:** RV failure is reported to involve mitochondrial dysfunction which might impair RV contraction by decreasing cardiomyocyte ATP level. Using the pulmonary artery constriction model of RV failure, we found that chronic treatment with an α1A-adrenergic receptor agonist increased RV myocardial respiration rate, increased RV myocardial ATP level, and increased RV function. These findings suggest that the α1A-adrenergic receptor is a therapeutic target for treating RV failure, and that the mechanism involves improved RV cardiomyocyte bioenergetic status.

## INTRODUCTION

Failure of the right ventricle (RV) occurs when the RV is unable to provide adequate blood flow through the pulmonary circulation at a normal preload (1). RV failure is a serious disease with a poor prognosis (1, 2). RV failure frequently arises in patients with failure of the left ventricle (LV) and causes markedly worse symptoms and prognosis compared to patients with LV failure without RV dysfunction (3, 4). Moreover, the prognosis of heart failure patients with pulmonary hypertension is strongly related to RV dysfunction (5). Unfortunately, pharmacologic therapies for RV failure are limited. Therapies developed to treat patients with LV failure are ineffective for improving function or survival of patients with RV failure (2). New therapies for RV failure are needed (6).

Previous studies suggest that powerful cardioprotective effects are mediated by α1-adrenergic receptors, in particular the α1A-subtype (α1A-AR) (7–11). In models of LV failure, chronic administration of an α1A-AR agonist protected against cardiomyopathy due to doxorubicin toxicity (9, 12), trans-aortic constriction (13) or myocardial infarction (10). The therapeutic effect was lost in α1A-AR knockout mice (9, 12, 13), suggesting that the therapeutic effect was mediated by α1A-AR signaling (rather than a non-specific drug effect). The therapeutic effect occurred at a low dose of α1A-AR agonist that did not raise blood pressure through vascular α1-AR activation (12). Moreover, we reported that this low therapeutic dose did not increase RV contraction in sham-operated animals, suggesting this dose did not stimulate an inotropic response in-vivo (14).

We reported that chronic α1A-AR agonism was beneficial in 2 models of RV failure: pulmonary fibrosis; and pulmonary artery constriction (14, 15). Chronic α1A-AR agonism resulted in less cardiomyocyte necrosis and greater RV function in-vivo (14, 15).

RV failure was recently reported to involve impaired mitochondrial respiration evidenced by impaired oxygen flux measured in RV homogenate and by increased production of reactive oxygen species (ROS), suggesting mitochondrial damage (16). In heart failure, impaired mitochondrial function resulting in impaired bioenergetic status could directly lead to contractile dysfunction (17). Indeed, we reported that reduced ATP level impaired the contraction of RV myocardium in-vitro (18). Thus, impaired mitochondrial function may play a role in RV failure and improved mitochondrial function might contribute to reversal of RV failure.

The goal of this study was to determine if reversal of RV failure after chronic treatment with an α1A-AR agonist involved increased mitochondrial function. We found that RV failure induced by pulmonary artery constriction (PAC) resulted in decreased tissue ATP level measured in RV homogenate, and decreased oxygen consumption rate by RV homogenate, which together indicate impaired mitochondrial function. Two wk after PAC, chronic treatment for a further 2 wk with the α1A-AR agonist A61603 resulted in significant recovery of RV function, and both increased ATP level and increased oxygen consumption rate measured in RV homogenates. These findings suggest that reversal of RV failure by chronic treatment with α1A-AR agonist involved increased mitochondrial function.

The current study confirms our previous report that RV failure is reversed by chronic treatment with α1A-AR agonist, demonstrating a robust finding (14). Together, our studies suggest that the α1A-AR is a novel therapeutic target to treat RV failure.

## METHODS

This institution is accredited by the American Association for the Accreditation of Laboratory Animal Care (Institutional PHS Assurance Number is A3476-01). The study was approved by the Animal Care and Use Subcommittee of the San Francisco Veterans Affairs Medical Center and conformed to the *Guide for the Care and Use of Laboratory Animals* published by the National Institutes of Health (Revised 2011).

### RV failure model and A61603 therapy

Adult male C57BL/6J mice (Jackson Laboratory) were used, age ≈ 11 weeks, body weight ≈ 26 g, at the beginning of the experiment. We used a pulmonary artery constriction (PAC) model of RV failure (RVF) as previously described (19). Animals were anesthetized with isoflurane (3% induction, 1.5% maintenance), intubated, and a left lateral thoracotomy performed. A 7-0 suture was placed around the pulmonary artery and used to tie the artery to a 26-gauge needle. The needle was quickly removed resulting in a ligature constricting the pulmonary artery. The chest and skin were sutured closed. Sham control surgeries were identical, except that the suture around the artery was not tied.

For the treatment group, 2 wk after PAC, mice were chronically treated for a further 2 wk with A61603 (Tocris Biotechne, Minneapolis, MN) a potent and highly specific agonist for the α1A-AR that does not stimulate the other two α1-AR-subtypes (α1B or α1D). A low dose of A61603 (10 ng/kg/day) (12) was given by continuous subcutaneous infusion with an osmotic mini-pump (Alzet, model #1002, Durect) implanted between the scapulae under isoflurane anesthesia. This dose was reported not to increase systolic or diastolic blood pressure, and to be close to the EC_50_ dose for activation of cardiac ERK signaling (12). For the vehicle group, 2 wk after PAC, mice were implanted with mini-pumps delivering only saline for a further 2 wk.

### RV Function

Fractional shortening of the RV outflow tract (RVOT FS) was measured using a Prospect T1 High Frequency Ultrasound Imaging System for Mice with 40 MHz Probe according to a published procedure (20).

### ATP level

ATP level in myocardial homogenate was measured using a luciferase assay (21). Hearts were flushed with PBS, RV and LV dissected and flash frozen. Frozen tissue was placed in Tris–EDTA–saturated phenol (Phenol-TE Sigma P-4557) and homogenized. Homogenate was transferred into a microtube containing Chloroform (Sigma c-2432) and de-ionized water, thoroughly shaken for 20 s, and centrifuged at 10,000×g for 5 min at 4°C. 200ul of diluted supernatant was used for ATP measurement by the CellTiter-Glo luminescent cell viability assay (Promega) as per manufacturer’s instructions.

### Respirometry

Punch biopsy samples (2 mm diameter, 1.5-3 mg wet wt) from the RV free wall were homogenized using an ice cold PBI-Shredder motorized tissue homogenizer with 0.5 mL of ice cold MiR05 respiration buffer (Oroboros Instruments, Innsbruck, Austria) and then diluted into a final volume of 5 mL and kept on ice until measurement. An Oroboros O2k High-Resolution Respirometer (Oroboros Instruments, Innsbruck, Austria) was used to measure O_2_ consumption by heart homogenates. The tissue homogenate was added into the two 2-mL glass chambers in the respirometer with stirring set at 750 RPM. Catalase (280 U/mL) was added to the mixture and O_2_ level was increased to about 450 µM by addition of H_2_O_2_. We measured, in sequence: O_2_ consumption linked to CI after addition of 10 mM glutamate, 5 mM pyruvate, 2.5 mM malate, and 5 mM ADP with 3 mM MgCl_2_; maximum CI- and CII-linked uncoupled respiration after addition of 10 mM succinate and 1.5 µM carbonyl cyanide m-chlorophenylhydrazone (CCCP, protonophore); and respiration linked to CII after inhibiting CI with 0.5 µM rotenone (22). Non-mitochondrial respiration was measured at the end of the protocol by inhibiting CII with 2.5 µM antimycin A and subtracted from all measurements.

### Histology and Electron Microscopy

Myocardial samples were immobilized by pinning, fixed overnight in buffered 2.5% glutaraldehyde, rinsed, osmicated, dehydrated and embedded in EPON. Ultrathin transverse sections were stained with toluidine blue or lead citrate/uranyl acetate using standard methodology.

### Western Blot Analysis

Snap-frozen RV free-wall tissue was homogenized in 200 μl ice-cold radioimmunoprecipitation (RIPA) buffer (150 mmol/L NaCl, 50 mmol/L Tris HCl pH 7.4, 1% NP-40, 0.5% sodium deoxycholate, 0.1% SDS, protease and phosphatase inhibitors) using plastic pestle in 1.5 ml Eppendorf tube. The lysate was snap frozen in liquid nitrogen and stored at −80°C. Lysate was thawed on ice and centrifuged at 1000 x G for 10 min to remove insoluble material. Protein concentration was measured using DC protein assay (Bio-Rad) and equilibrated to 35 μg of protein in each sample. Proteins were separated by SDS-PAGE on 12% gels (Bio-Rad) and transferred to a nitrocellulose membrane. Proteins were detected with the following primary antibodies: acetylated lysine (Invitrogen MA1-2021), Citrate Synthase (Invitrogen GT1761); phospho-p44/42 MAPK (p-ERK) (Cell Signaling #4370), p44/42 MAPK (Total ERK) (Cell Signaling #9102); 4-hydroxynonenal (4-HNE, Abcam Ab46545); Glutathione Peroxidase-1 (Biotechne AF3798). Bands were developed using SuperSignal West Dura ECL reagent (ThermoFisher), captured with the ChemiDoc XRS camera system and analyzed with QuantityOne software version 4.6.9. Indian Ink staining of the quantified blots confirmed equal loading of protein in each lane.

### Statistical analysis

Data are presented as mean ± SE. Statistical tests (paired or unpaired t-test, and 1-way or 2-way ANOVA with post hoc analysis using Dunnett’s test for multiple comparisons) were performed using Prism software (GraphPad Software, Inc., La Jolla, CA) with a significance level set at *P*<0.05.

## RESULTS

### Chronic α1A-AR agonist treatment reversed RV failure

We used echocardiography to assess RV function from the fractional shortening of the RV outflow tract (RVOT FS) (23). Prior to agonist treatment, RVOT FS in sham (48.7 ± 0.8%, n = 9) was significantly decreased 2 wk after pulmonary artery constriction (PAC) (30.1 ± 0.9 %, n = 27, *P* <0.001, unpaired t test) (Fig. 1A). After a further 2 weeks, the RVOT FS for sham controls (46.5 ± 1.3 %, n = 9) was reduced with vehicle treatment (27.6 ± 1.5 %, n = 13, *P* < 0.0001), but recovered with A61603 treatment (34.5 ± 0.6 %, n = 14, *P* < 0.001) (Fig. 1B). Relative to pre-treatment values, RVOT FS was increased after chronic A61603 treatment (20.2 ± 2.6%, n = 14), but not after vehicle treatment (−11.5 ± 3.5%, n = 13, *P* < 0.0001) (Fig. 1C).

**Figure 1.**
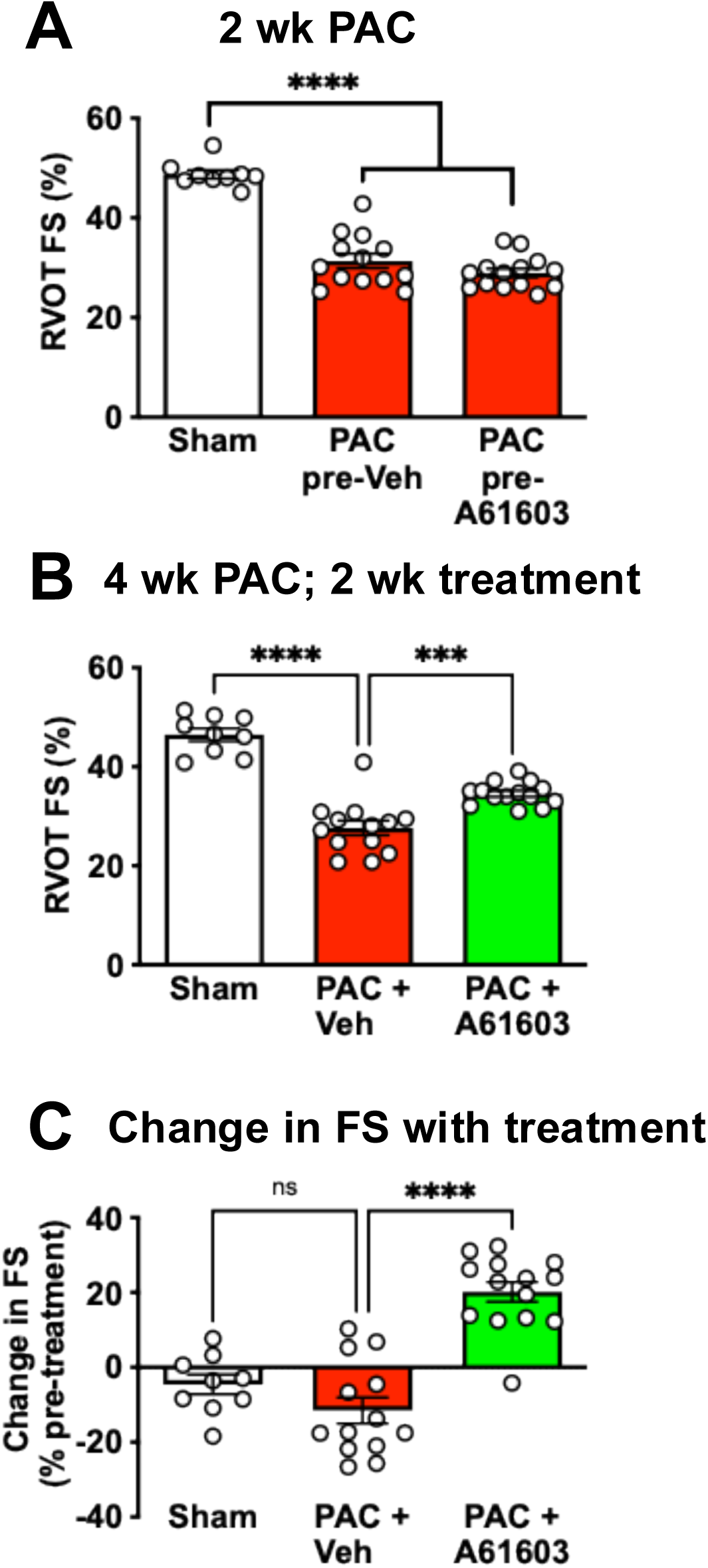
Chronic A61603 treatment improved RV function assessed as RV outflow tract fractional shortening (RVOT FS) using echocardiography. (A) 2 wk after surgery and prior to treatment, RVOT FS was reduced after PAC (combined n = 27) compared to sham-operated (n = 9) (**** P < 0.0001, unpaired t test). (B) 4 wk after PAC, RVOT FS was higher after 2 wk treatment with A61603 (10 ng/kg/day) vs. vehicle treatment (**** P < 0.0001, *** P < 0.001, one-way ANOVA with Dunnet post-hoc test). (C) the increase in RVOT FS (before vs. after treatment) was highly significant (**** P < 0.0001, ns = not significant, one-way ANOVA with Dunnet post-hoc test).

Consistent with these functional measures, the ratio of liver weight to body weight for the sham group (41.6 ± 1 mg/g, n = 22) was increased 4 wk after PAC plus vehicle treatment, consistent with liver congestion, a characteristic symptom of RV failure (47.1 ± 0.7 mg/g, n = 28, *P* < 0.0001) (Fig. 2A). However, 4 wk after PAC with 2 wk chronic treatment with A61603, the ratio of liver weight to body weight was lower (43 ± 0.7 mg/g, n = 26, *P* < 0.001), consistent with reversal of RV failure.

**Figure 2.**
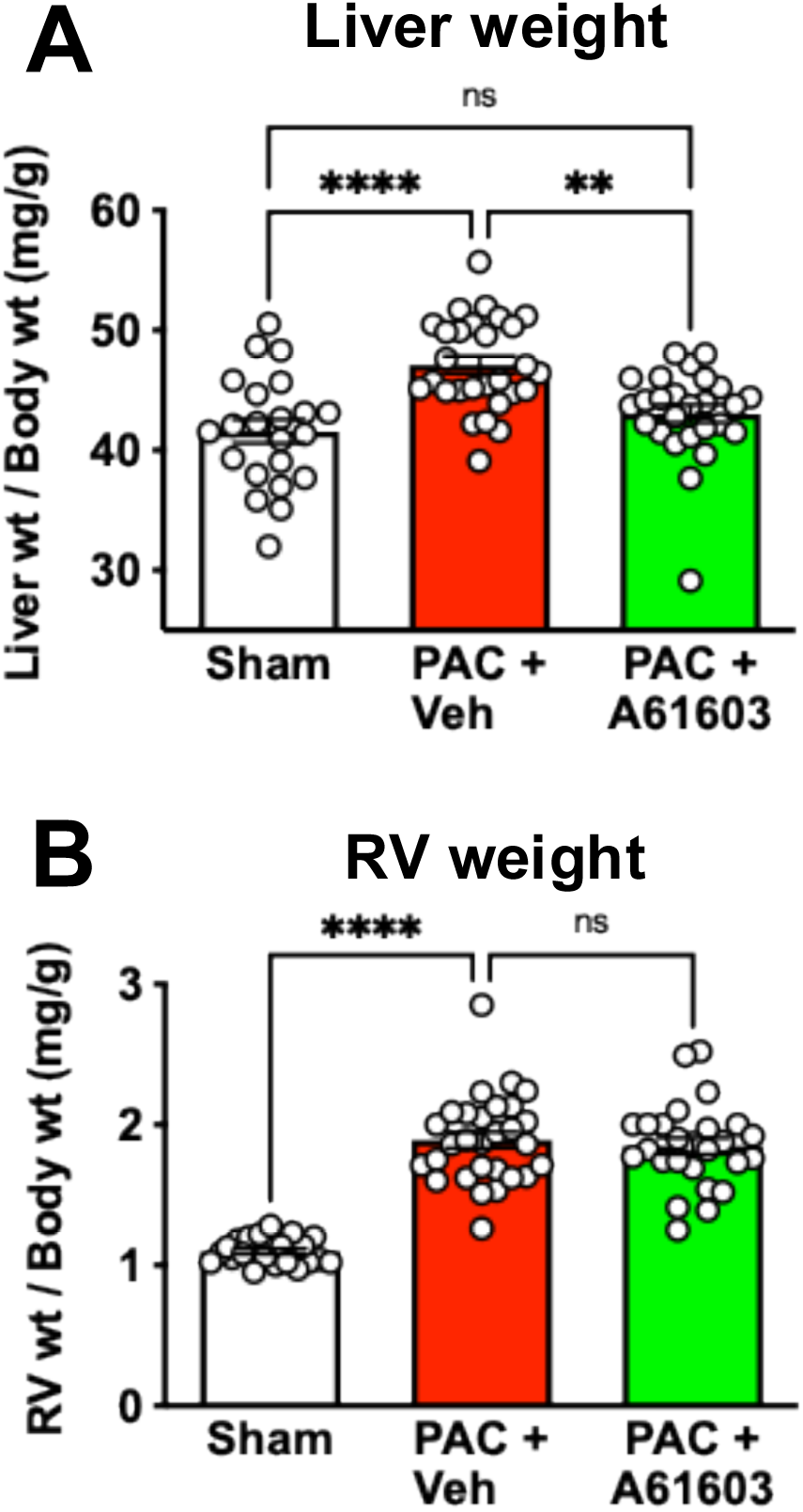
Morphological effects of PAC and chronic A61603 treatment. 4 wk after PAC, 2 wk treatment with A61603 reduced liver congestion (lower liver weight) **(A)**, but did not reduce RV hypertrophy (RV weight) **(B)**. (**** P < 0.0001, ** P < 0.01, ns = not significant, one-way ANOVA with Dunnet post-hoc test).

RV pressure overload due to PAC was associated with marked RV hypertrophy. The ratio of RV weight to body weight in sham (1.1 ± 0.02 mg/g, n = 22,) was appreciably increased 4 wk after PAC with 2 wk vehicle treatment (1.9 ± 0.06 mg/g, n = 29, *P* < 0.0001) (Fig. 2B). RV hypertrophy after PAC was not reduced after 2 wk treatment with A61603 (1.8 ± 0.06 mg/g, n = 26, *P* = 0.8) (Fig. 2B), consistent with our previous report (14).

### Chronic α1A-AR agonist treatment increased RV tissue bioenergetic status

We measured the ATP level in RV tissue homogenates using a luciferase assay. Compared to Sham controls, tissue homogenate ATP level (3.3 ± 0.1 mM, n = 10) was reduced by 43% 4 wk after PAC with 2 wk vehicle treatment (1.9 ± 0.1 mM, n = 6, *P* < 0.0001) (Fig. 3A). Importantly, 4 wk after PAC, ATP level in RV tissue homogenates was appreciably higher for animals that were treated for 2 wk with A61603 (2.6 ± 0.13 mM, n = 7, *P* < 0.01) (Fig. 3A). Thus, chronic A61603 treatment had a beneficial effect on bioenergetic status. Previously, we reported that PAC resulted in RV tissue fibrosis (≈ 20%) that was not reversed by chronic A61603 treatment (14). Thus, the increased ATP level we observed after chronic A61603 treatment was likely not due to decreased tissue fibrosis.

**Figure 3.**
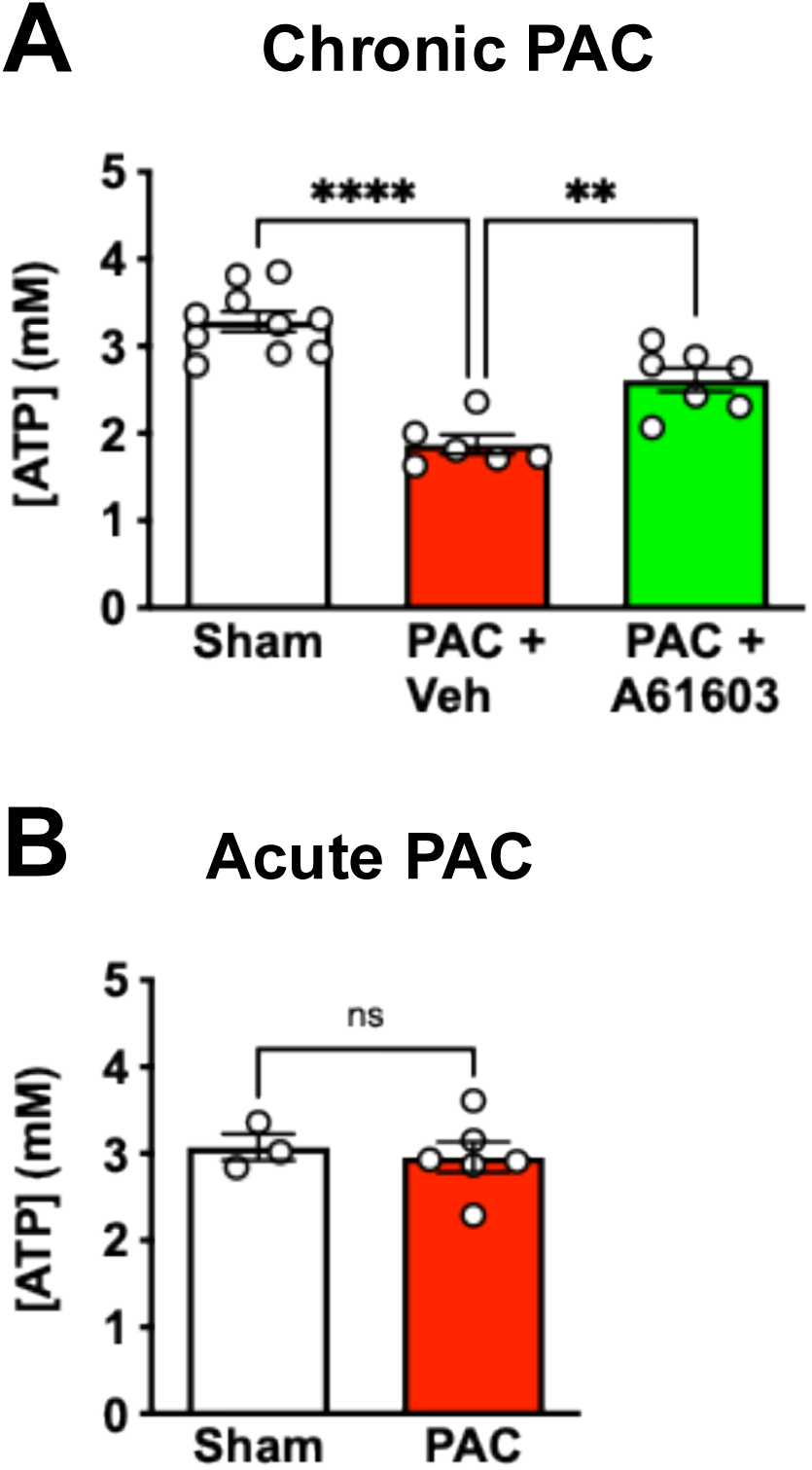
Chronic A61603 treatment increased tissue ATP level. **(A)** 4 wk after PAC, 2 wk treatment with A61603 increased tissue ATP level (**** P < 0.0001, ** P < 0.01, one-way ANOVA with Dunnet post-hoc test). (**B**) Acute PAC for 30 min. did not reduce ATP level (ns = not significant, unpaired t test).

In additional control experiments, we found that the ATP level in RV tissue homogenate was not reduced by a brief (30 min.) episode of PAC (3.0 ± 0.18 mM, n = 6) compared to Sham (3.1 ± 0.15 mM, n = 3, *P* > 0.05, unpaired t test) (Fig. 3B). Thus, PAC did not directly reduce ATP level acutely, for example, due to myocardial ischemia.

To further probe bioenergetic metabolism, we measured mitochondrial respiration (O_2_ consumption) fueled by substrates linked to Complex I (CI), or Complex II (II), as well as the maximum respiration rate of uncoupled mitochondria.

For RV tissue homogenates, the mitochondrial respiration rate (represented by O_2_ flux) was reduced after PAC (Fig. 4), consistent with previous studies reporting that RV failure involved impaired mitochondrial function (16, 22, 23). For CI-linked respiration, O_2_ flux observed in sham (362 ± 34 pM/s/mg, n = 9) was reduced by 41% at 4 wk after PAC plus vehicle treatment (216 ± 17 pM/s/mg, n = 15, *P* < 0.001) (Fig. 4A). RV homogenates from animals 4 wk after PAC with 2 wk A61603 treatment had higher O_2_ flux (294 ± 20 pM/s/mg, n = 11, *P* < 0.05) (Fig. 4A).

**Figure 4.**
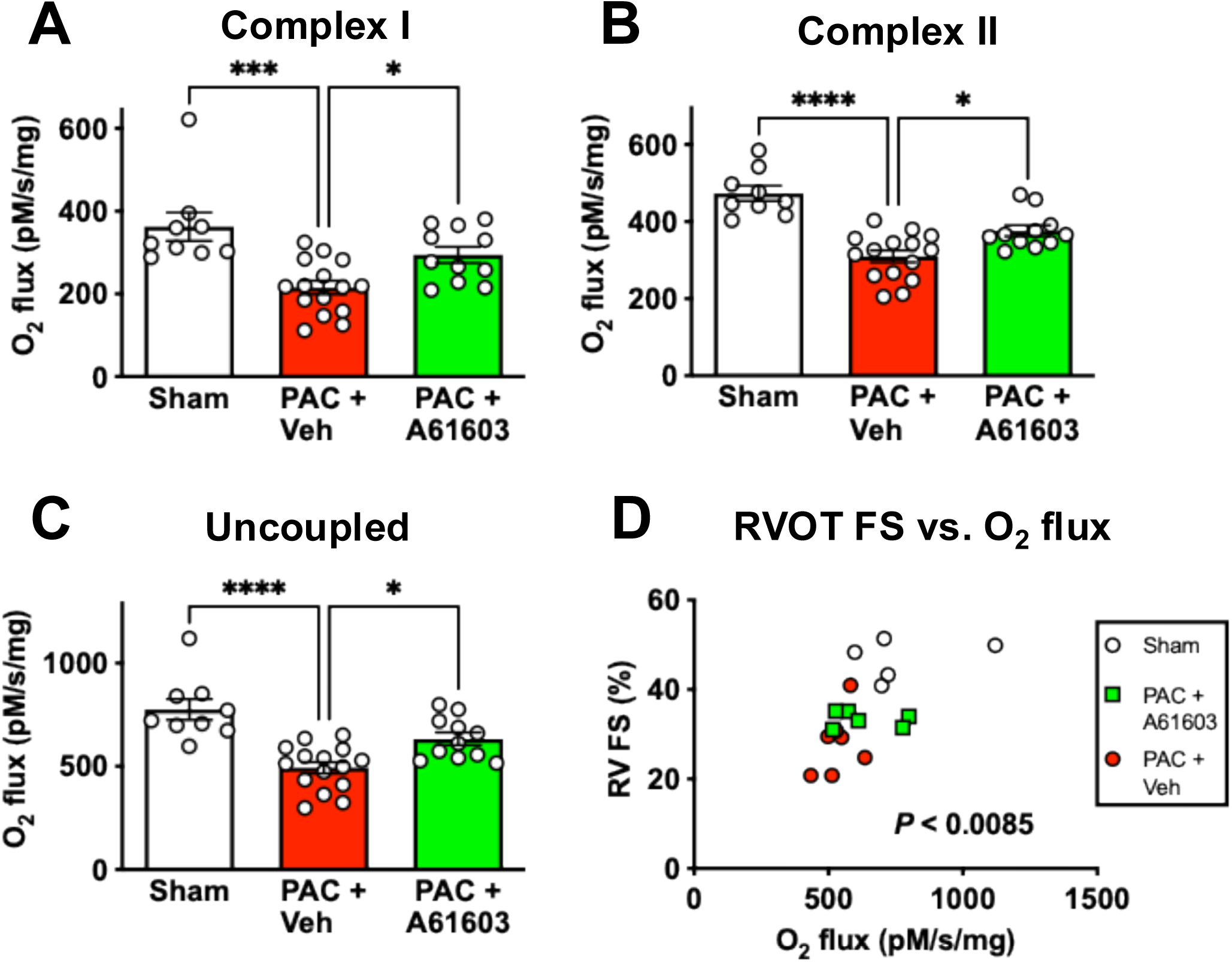
Chronic A61603 treatment increased tissue respiration. 4 wk after PAC, 2 wk treatment with A61603 increased tissue respiration rate driven by Complex I (**A**), Complex II (**B**), and maximum uncoupled respiration (C). There was a significant relation between RV function and maximum respiration rate (D). (**** P < 0.0001, *** P < 0.001, * P < 0.05, one-way ANOVA with Dunnet post-hoc test).

The effect of chronic A61603 treatment to increase mitochondrial respiration rate likely contributed to the increased ATP level observed after chronic A61603 treatment. Decreased respiration rate after PAC and recovery of respiration rate after chronic A61603 treatment were also observed in measures of respiration linked to CII (Fig. 4B) and for maximum uncoupled respiration (Fig. 4C).

Consistent with the respiration rate having functional importance, there was a significant relationship between RV function (RVOT FS) versus the maximum uncoupled respiration rate (Fig. 4D). Thus, impaired respiration rate after PAC might contribute to decreased RV function and the effect of chronic treatment with A61603 to increase respiration rate might contribute to recovery of RV function.

To determine if the recovery of mitochondrial respiration rate after chronic A61603 treatment involved increased mitochondrial content per cell we used electron microscopy to estimate the percent area of the myocyte that was occupied by mitochondria. The cell area occupied by mitochondria for the sham group (33.3 ± 0.3%, n = 4) was reduced 4 wk after PAC with vehicle treatment (25.6 ± 2.4%, n =5, *P* < 0.01) (Fig. 5A). But 2 wk A61603 treatment did not increase the cell area occupied by mitochondria (21.6 ± 0.9, *P* > 0.05). Thus, a decrease in mitochondrial content after PAC may have contributed to the decreased bioenergetic status after PAC. However, the recovery of bioenergetic status observed after PAC with chronic A61603 treatment did not involve an increase in the mitochondrial content per cell.

**Figure 5.**
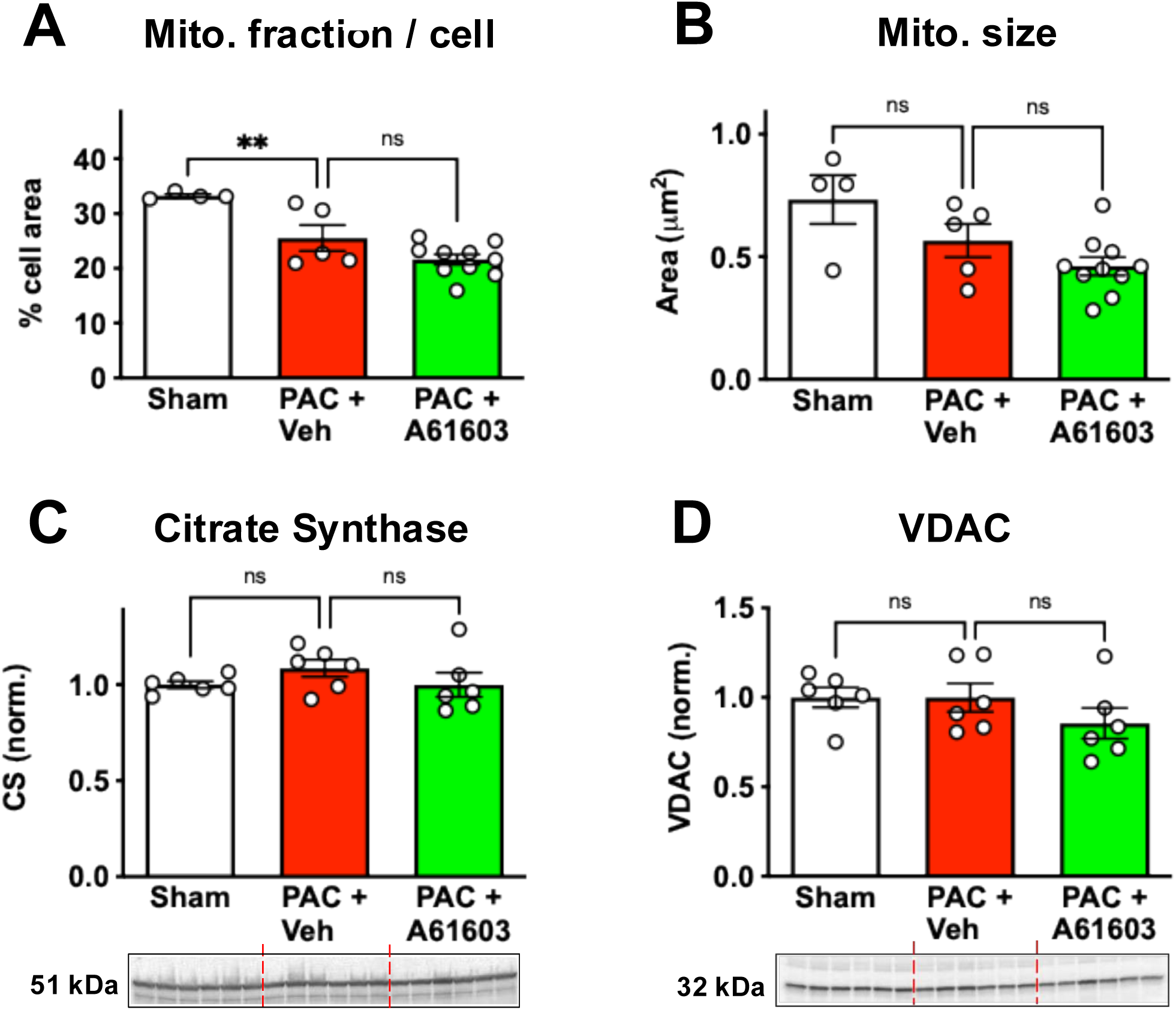
Chronic A61603 treatment did not increase mitochondria abundance. Quantitation of EM micrographs show that chronic A61603 did not increase the cell area occupied by mitochondria (**A**), or mitochondria size (**B**). Western blots show that chronic A61603 treatment did not increase the abundance of citrate synthase or VDAC (**D** & **E**). (** P < 0.01, ns = not significant, one-way ANOVA with Dunnet post-hoc test).

The average area per individual mitochondrion for the sham group (0.73 ± 0.1 μm^2^, n = 4) trended smaller 4 wk after PAC but not significantly (0.57 ± 0.07 μm^2^, n = 5, *P* > 0.05) (Fig. 5B). After PAC, there was not an increase in mitochondria size that occurred after 2 wk. treatment with A61603 (0.46 ± 0.04 μm^2^, n = 10, *P* > 0.05) (Fig. 5B).

Finally, despite the reduction in cell area occupied by mitochondria after PAC, levels of the mitochondrial proteins citrate synthase and VDAC (voltage-dependent anion channel) were not decreased after PAC plus vehicle treatment, and were not affected by chronic A61603 treatment (Figs. 5C, 5D).

Together, these results suggest that the recovery of mitochondrial function after A61603 treatment did not involve an increase in mitochondrial size or abundance.

### Chronic α1A-AR agonist treatment induced cardioprotective signaling

Activation of the pro-survival signaling kinase ERK was assessed from the ratio of phosphorylated ERK to total ERK. When normalized to sham controls, we found that 4 wk after PAC, ERK activation was higher in animals treated for 2 wk with A61603 (1.56 ± 0.36, n = 6) compared to vehicle-treated (0.56 ± 0.11, n = 6, *P* < 0.05) (Fig. 6A), consistent with our previous report (14).

**Figure 6.**
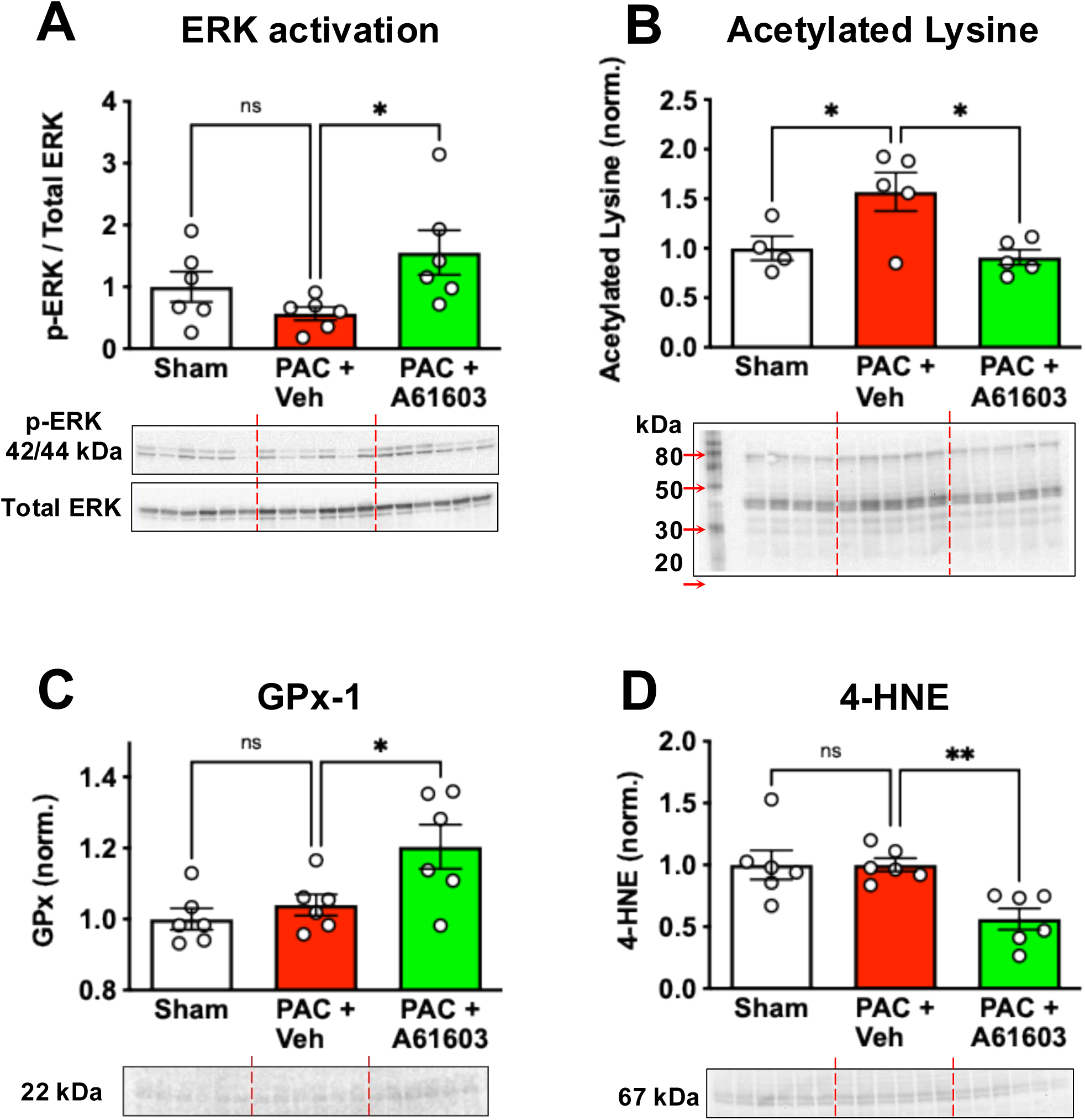
Chronic A61603 treatment stimulated cardioprotective effects. 4 wk after PAC, 2 wk treatment with A61603 increased ERK signaling (**A**), decreased acetylated lysine level (**B**), increased glutathione peroxidase-1 (**C**) and decreased the oxidation product 4-HNE (**D**). (** P < 0.01, * P < 0.05, ns = not significant, one-way ANOVA with Dunnet post-hoc test).

Protein hyperacetylation has been linked to mitochondrial dysfunction in heart failure (24–26). When normalized to sham controls, we found that 4 wk after PAC with 2 wk vehicle treatment there was protein hyperacetylation (1.57 ± 0.19, n = 5, *P* < 0.05) (Fig. 6B). However, protein acetylation level was not increased 4 wk after PAC with 2 wk. A61603 treatment (0.91 ± 0.08, n = 5, *P* < 0.05) (Fig. 6B).

RV myocardium 4 wk after PAC with 2 wk A61603 treatment had increased expression of the antioxidant GPx1 (1.2 ± 0.06, n = 6) vs. vehicle treatment (1.04 ± 0.03, n = 6, *P* < 0.05) (Fig. 6C), similar to our previous report (14). Consistent with increased antioxidant, the level of 4-HNE (lipid peroxidation product generated during oxidative stress) was reduced after PAC with 2 wk A61603 treatment (0.56 ± 0.09, n = 6) vs vehicle treatment (1.0 ± 0.05, n = 6, *P* < 0.01) (Fig. 6D), similar to our previous report (15).

## DISCUSSION

The overall finding of this study is that chronic treatment of the failing RV with the α1A-AR agonist A61603 rescued the bioenergetic status of failing RV myocardium. Reversing the decline in RV tissue bioenergetic status may contribute to the reversal of RV dysfunction that we observed after chronic α1A-AR-agonist treatment.

### Reversal of RV failure

In the setting of RV failure, the current study found that chronic α1A-AR agonism results in reversal of RV failure as evidenced by recovery of RV function in-vivo and decreased liver congestion. The current study confirms our previous study (14), and thus demonstrates that reversal of RV failure after chronic treatment with an α1A-AR agonist is a robust finding.

Currently, there is not an effective pharmacologic treatment for RV failure. Consistent with our previous reports, this study suggests that the α1A-AR is a novel therapeutic target for treating RV failure (14, 15, 27).

### Rescue of RV tissue bioenergetic status

In the setting of RV failure we found a reduced level of ATP and a reduced respiration rate that were measured in RV tissue homogenates. These findings are consistent with mitochondrial dysfunction that was reported to occur in the mouse PAC model of pressure overload-induced RV failure (16).

Relative to sham controls, we found that RV myocardial [ATP] level was reduced by 43% after PAC. The finding of reduced ATP level in RV failure is similar to the reduced ATP level reported in failing LV myocardium (28, 29). Previous studies suggested that the reduced cellular [ATP] in heart failure leads directly to inhibition of contraction (17). However, the actomyosin ATPase has a very high affinity for ATP (Km ATP 10-150 μM) (30, 31), and thus, may not be sensitive to a 50% fall of [ATP] that still remains within the mM range. Recently, we tested the effect of millimolar ATP levels on the contraction of RV myocardium. ATP levels were decreased within the range of 8 – 2 mM referenced to cytosolic water (18). We reported that a 50% reduction of ATP, to a level typically found in heart failure, caused appreciably reduced mechanical power generation and slowed cross-bridge kinetics (18). This suggests that the magnitude of the decrease in [ATP] that we observed in RV failure could contribute to impaired RV function.

The current study found that chronic A61603 treatment appreciably increased RV myocardial ATP level. Based on our report that myocardial contraction is influenced by ATP level in the millimolar range (18), the increased ATP level resulting from chronic A61603 treatment might contribute to the recovery of RV function that we observed.

Recovery of ATP level after chronic A61603 treatment was closely linked to increased mitochondrial respiration measured in RV homogenate. There was not an increased area fraction of mitochondria in myocytes after chronic A61603 treatment. The abundance of the mitochondrial markers citrate synthase and VDAC were not increased after chronic A61603 treatment. Finally, regression of fibrosis was not observed after chronic A61603 treatment (14). Together these results suggest that the increased respiration rate observed was not due to increased abundance of mitochondria but likely involved increased mitochondrial intrinsic function.

We observed several molecular signatures that were reported to be associated with increased mitochondrial function. Consistent with our previous report, we found that chronic A61603 treatment resulted in activation of the signaling kinase ERK (14). An α1A-AR – ERK survival signaling pathway mediates increased survival in-vitro of cardiomyocytes challenged with norepinephrine, doxorubicin or H_2_O_2_, and protection in-vivo against doxorubicin cardiotoxicity (9, 12, 32). Pro-survival effects of ERK signaling were associated with increased mitochondrial membrane potential, inhibition of mitochondrial permeability transition pore opening, increased mitochondrial ATP synthesis, and decreased mitochondrial oxidative stress (33–35).

We found lysine hyperacetylation after PAC, a post-translational modification previously associated with decreased mitochondrial function and heart failure (24, 25, 36). In the PAC model of RV failure, we found that lysine hyperacetylation was reversed after 2 wk treatment with A61603, which may have contributed to the recovery of mitochondrial function we observed.

Finally, consistent with our previous reports, we found that chronic treatment with A61603 was linked to decreased oxidative stress, evidenced by increased levels of the antioxidant GPx-1 and decreased oxidant product 4-HNE (14, 15). Oxidative stress damages multiple mitochondrial components resulting in decreased mitochondrial function (8, 37, 38). Decreased oxidative stress after chronic treatment with A61603 might contribute to increased mitochondrial function.

Our findings in RV failure, are consistent with multiple studies by Jensen and coworkers showing that chronic α1A-AR stimulation improved mitochondrial function in models of LV failure (9–11). In a doxorubicin cardiomyopathy model, chronic treatment with the α1A-AR agonist dabuzalgron preserved ATP content in-vivo and rescued mitochondrial function in-vitro (9). Following myocardial infarction, chronic treatment with α1A-AR agonist A61603 protected against oxidative stress and increased mitochondrial Complex I activity (10). In contrast, knockout mice lacking α1A-ARs in the heart have higher mortality after myocardial infarction (11).

### Limitations

Multiple therapeutic effects linked to chronic α1A-AR activation have been described, including decreased necrosis, decreased oxidative stress, increased myofilament contraction, and increased ATP level (7). However, the relative contribution of these effects and the detailed underlying mechanisms remain unclear.

We previously reported that the α1A-AR was present in 60% of myocytes (39). In the current study, it is not known if the improved mitochondrial function observed involved only a subset of RV cardiomyocytes.

## Conclusion

Chronic treatment of RV failure with α1A-AR agonist mediates reversal of RV failure involving increased tissue ATP level and increased mitochondrial function. The current study is consistent with our previous suggestion that the α1A-AR is an effective therapeutic target to reverse RV failure.

## ACKNOWLEDGEMENTS

This work was supported by Department of Veterans Affairs Merit Review Awards I01BX000740 and I21BX006858 (AJB); and National Heart, Lung and Blood Institute Grant R01 HL154624 (AJB). Expert technical assistance of Guanying Wang is gratefully acknowledged.

